# dandelionR: Single-cell immune repertoire trajectory analysis in R

**DOI:** 10.1101/2025.02.04.636146

**Authors:** Jiawei Yu, Xiaohan Xu, Nicholas Borcherding, Zewen Kelvin Tuong

## Abstract

Integration of single-cell RNA-sequencing (scRNA-seq) and adaptive immune receptor (AIR) sequencing (scVDJ-seq) is extremely powerful in studying lymphocyte development. A python-based package, *Dandelion*, introduced VDJ-feature space method, which addresses the challenge in integration of single-cell AIR data with gene expression data and enhanced the trajectory analysis results. However, no R-based equivalent or similar methods currently exist. To fill this gap, we present *dandelionR*, an R implementation of *Dandelion*’s trajectory analysis workflow, bringing the VDJ feature space construction and trajectory analysis using diffusion map and absorbing Markov chain to R, offering a new option for scRNA-seq and scVDJ-seq analysis to R language users.

## Introduction

During the development of T- and B-cells, variable (V), diversity (D) and joining (J) genes of adaptive immune receptors (AIR) recombine stochastically, introducing variability in the joining region (Davis & Bjorkman, 1988; Tonegawa, 1983). This process, known as V(D)J recombination, plays a critical role in generating diversity of the AIR repertoire (AIRR) (Carmona & Schatz, 2017). The diversity of AIRR is essential to adaptive immunity (Scott & Breden, 2020), but remains challenging for single-cell RNA sequencing (scRNA-seq) analysis (Heumos et al., 2023). This challenge necessitates the integration of scRNA-seq with adaptive immune receptor (AIR) sequencing (scVDJ-seq) (Irac et al., 2024).

*Scirpy* (Sturm et al., 2020), *Dandelion* (Stephenson et al., 2021; Suo et al., 2024), and *scRepertoire* (Borcherding et al., 2020; Yang et al., 2024) are widely used tools for conducting scVDJ-seq analysis. Among these, *Dandelion*’s Python-based innovative strategy of creating a VDJ feature space addressed some challenges in integrating AIR data within scRNA-seq, arising from the mixture of categorical and continuous data characteristics inherent to AIRR data. The feature space was leveraged to enable trajectory analysis informed by both the gene expression and VDJ data, which improved the prediction accuracy of trajectories from double-positive T cells to CD4/CD8 T cells, demonstrating significant potential for future applications (Suo et al., 2024).

Here, we introduce *dandelionR*, an R-based scVDJ-seq trajectory analysis tool replicating the trajectory analysis workflow of *Dandelion. dandelionR* enables the construction of the VDJ feature space to perform trajectory analysis using diffusion maps and absorbing Markov chains, with seamless interaction with *scRepertoire*. The current version (v0.99.0) is available on GitHub (https://www.github.com/tuonglab/dandelionR) and under review at Bioconductor, along with user documentation and additional resources. This tool addresses existing gaps in functionality among current tools, offering researchers a more convenient solution for analysing immune repertoires and single-cell sequencing data in R. By doing so, it facilitates a deeper exploration of lymphocyte development and its functional mechanisms.

## Methods

For trajectory analysis, *Dandelion* requires cell pseudobulks, typically with *Milo* (Dann et al., 2022), to construct the pseudobulked VDJ feature space. The feature space is then used as the input for *Palantir* (Setty et al., 2019), a trajectory analysis tool which employs diffusion maps and absorbing Markov chain to infer trajectory. *Palantir* produces pseudotime values and probabilities of each pseudobulk, which *Dandelion* subsequently projects back to each cell.

*dandelionR* is developed and tested in R v4.4.1 to integrate with the Bioconductor framework and can interact with *scRepertoire* v2.2.1 onwards. As an R implementation of *Dandelion*, it aims to reproduce the pre-processing, feature-space-building and result-projecting functions of the original software.

The typical workflow of *dandelionR* proceeds as follows:

### Input

*dandelionR* uses a *SingleCellExperiment* object already combined with vdj data, such as from reading with *scRepertoire* (Borcherding et al., 2020; Yang et al., 2024), or processed using the python package *Dandelion* with *AnnData* and converted to *SingleCellExperiment*.

### Pre-processing

This step includes filtering cells with non-productive immune receptors and ambiguous VDJ chain status, e.g., orphan/incomplete or multiple TCRs in one cell, retaining only cells with relevant or complete VDJ data. Then, the remaining VDJ contigs that express the highest UMI counts in a cell are extracted for downstream analyses. Depending on the data source, some pre-processing steps may not be necessary and can be skipped or modified. For example, when using *scRepertoire*-derived data, the user should set ‘*already*.*productive = TRUE’* to skip the productive filtering process, as the filtering has already been handled as part of *scRepertoire*’s standard workflow. Additionally, there are many parameters that users can adjust according to their analysis requirements. For example, they can set ‘*allowed_chain_status = NULL*’ to skip checking whether a cell has relevant TCR chains and accept all contigs. This flexibility allows for a highly customizable pre-processing workflow as per user requirements. A default set of parameters has been defined based on the original *Dandelion* workflow to replicate the initial findings (Suo et al., 2024).

### Pseudobulking and feature space constructing

Pseudobulking can be achieved through the *miloR* (v2.0.0) package (Dann et al., 2022). Within each pseudobulk, *dandelionR* will tabulate the usage of each VDJ gene to create the VDJ feature space.

### Trajectory analysis

Using the constructed VDJ feature space as input, we can utilise trajectory analysis tools to obtain pseudotime and branching probabilities of each pseudobulk.

### Projection

The calculated pseudotime values and probabilities of each pseudobulk will be projected back onto individual cells, generating the final trajectory analysis results.

### Data

We converted the sample data from *Dandelion*’s tutorial, *demo-pseduobulk*.*h5ad*, into *SingleCellExperiment* format. The original data is from (Suo et al., 2022) and contains gene expression data with VDJ information.

## Results

### Replication of Workflow Before Trajectory Inference

To replicate the Dandelion workflow before trajectory analysis (**Fig. 1**), we implemented the following functions:

**Fig. 1.**
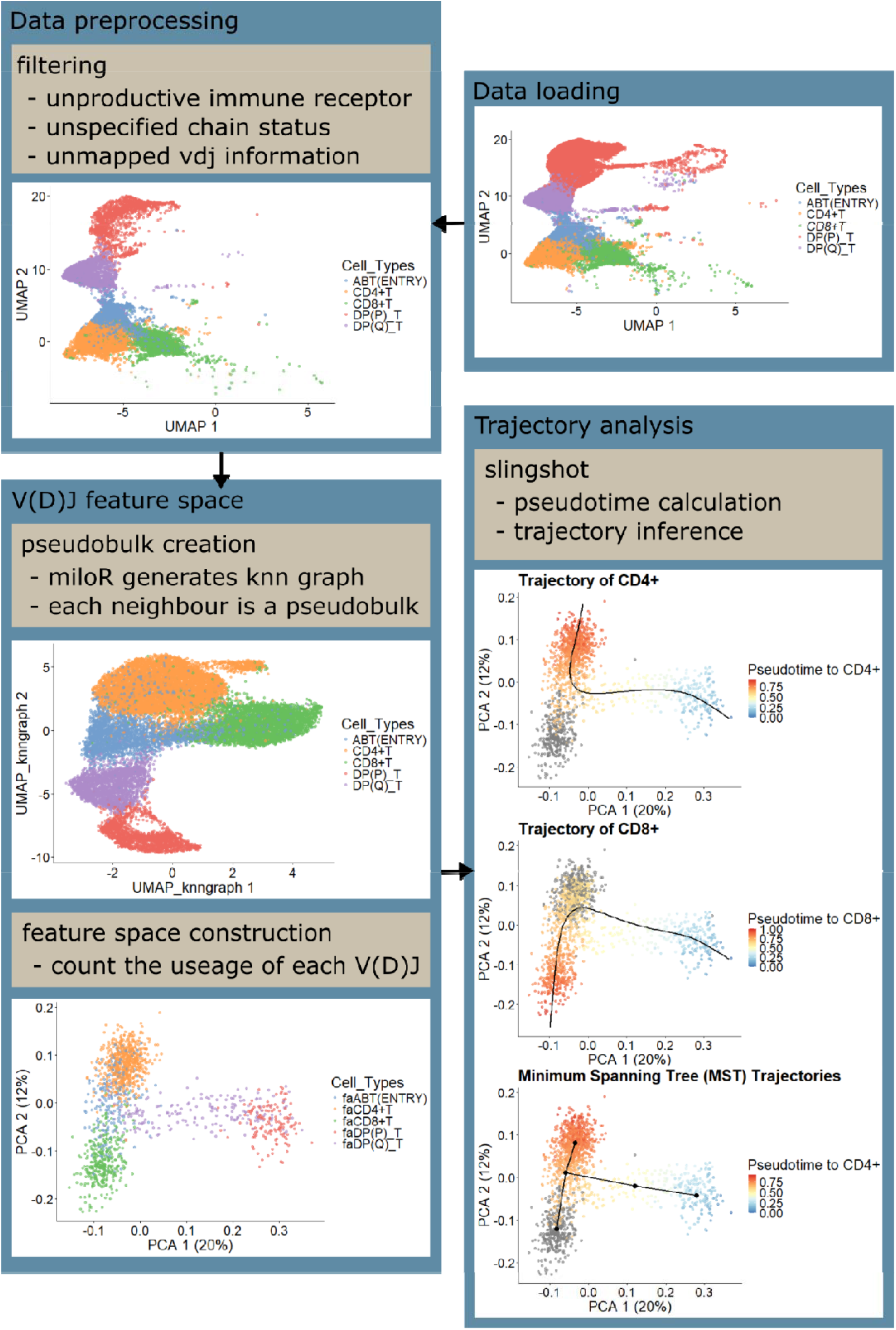
Overall workflow from pre-processing to trajectory analysis with *dandelionR*.

The *dandelionR::setupVdjPseudobulk* function pre-processed the demo single-cell VDJ data by Suo C *et al*. 2022. It then filtered out cells with non-productive or unclearly mapped alpha-beta TCR chains, extracting the main productive chain to store in a new column within the *colData* slot of the *SingleCellExperiment* object. Out of 65102 cells in the demonstration data, 17308 cells remained after filtering due to having complete TCR information necessary for downstream analyses.

Using *MiloR* (Dann et al., 2022), we constructed a k-nearest neighbour graph from the pre-processed data, treating each neighbour as a pseudobulk. This step allocated cells to pseudobulks based on the similarity of their gene expression profiles. Subsequently, the *dandelionR::miloUmap* function utilised the graph’s adjacent matrix to generate a UMAP (Uniform Manifold Approximation and Projection). The *dandelionR::vdjPseudobulk* function then created a VDJ feature space by counting the usage of each gene in each pseudobulk. With 160 V/D/J genes and 1516 pseudobulks, the VDJ feature space captured features from both gene expression and VDJ information in a continuous data format.

### Implementing trajectory inference based on absorbing Markov chains

The *Dandelion* workflow originally used the constructed feature space as an input for *Palantir*, a trajectory analysis tool, treating each pseudobulk as a cell and VDJ usage as gene expression information. *Palantir* employs probabilistic methods (Setty et al., 2019), which are primarily implemented in Python-based tools (Deconinck et al., 2021). However, most R-based tools do not incorporate such methods. Instead, *TSCAN* (Ji & Ji, 2016, 2019) utilises a self-developed travelling salesman problem (TSP) algorithm, *Slingshot* (Street et al., 2018) combines both minimum spanning tree with a self-modified principal curve, and *destiny* (Angerer et al., 2016) applies diffusion map. While *Ouija* (Campbell & Yau, 2019) is a probabilistic method utilising a Bayesian latent variable model, it is unsuitable for our dataset. This is because *Ouija* is limited to data with a linear topology, whereas we are certain that our dataset exhibits a bifurcation between CD4+ and CD8+ cells.

Since there are no direct *Palantir* equivalent or similar methods in R, we first used the trajectory analysis tools evaluation tool from *dynverse* (Saelens et al., 2019) to identify a suitable trajectory analysis tool for dandelion. We ran the function *dynguidelines::guidelines_shiny()*, setting the number of cells to 1516, the number of genes to 160, and the expected topology to bifurcation. Among all the evaluated tools, *Slingshot* stood out with top accuracy, followed by four Python-based tools.

We first attempted trajectory inference with *Slingshot* to obtain pseudotime and trajectory results (**Fig. 1**) (Street et al., 2018). *Slingshot* modifies the principal curve in two ways, one of which is incorporated with cell weights. This helps assign cells to lineages. Additionally, the common point of origin and weight function ensure that pseudotime values remain close before two lineages bifurcate. The author also recommended using the calculated weights to identify lineage-specific genes. These features suggest its potential to model cell fate.

To explore whether “cell weights” can serve as a substitute for branching probabilities in *Palantir*, we plotted pseudobulks coloured by weight (**Fig. 2**). Notably, cells located before a divergence typically receive weights close to 1 for both trajectories, indicating these cells should be assigned and have their pseudotime calculated within the lineage. However, for branching probability, we expected a value around 0.5, suggesting that the cell will differentiate to either a CD4+ T cell or a CD8+ T cell with similar probability. Additionally, the probability of differentiation into CD4^+^ T cells could be slightly higher, as CD4^+^ T cells are more abundant in an individual.

**Fig. 2.**
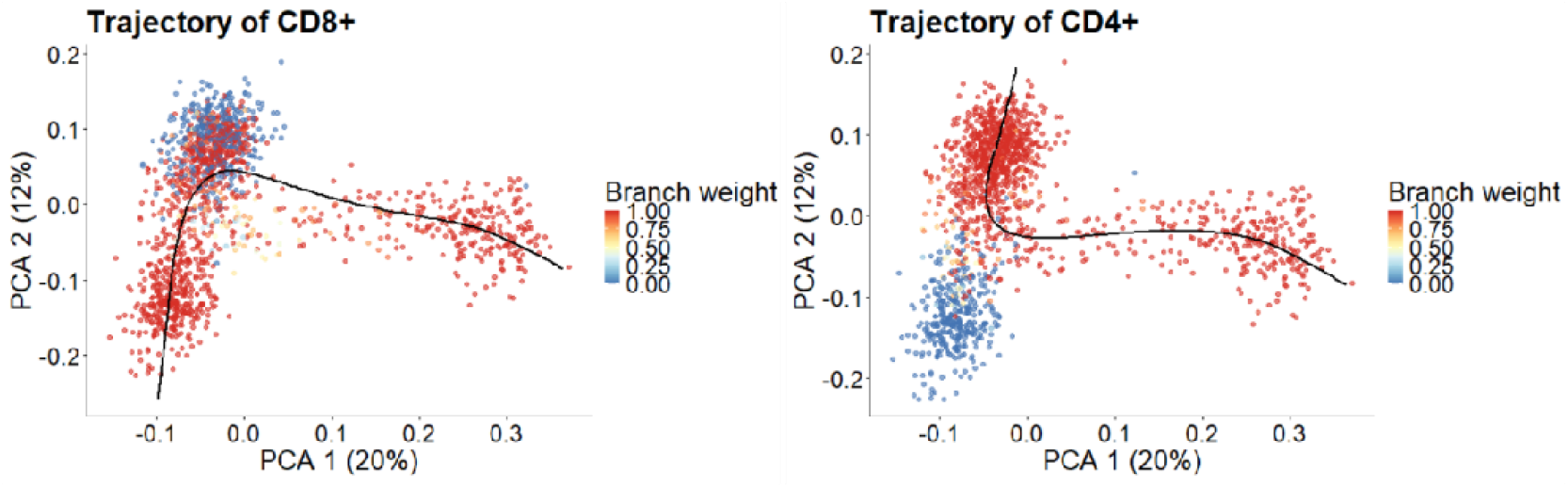
Weights produced by *Slingshot*. Each dot represents a single pseudobulk coloured according to the weight values assigned by *Slingshot* within each lineage curve. The colour palette scales from 0 (blue) to 1 (red), where 0 indicates that the data point does not belong to the respective lineage.

Since *Slingshot* could not produce values that substitute for branching probability, we sought to implement *Palantir*’s trajectory analysis function in R. We anticipate that this approach could not only address the lack of branching probability in our workflow but also help fill a critical gap in the R community, where probabilistic methods for trajectory analysis remain scarce.

In the original *Dandelion* workflow, *Palantir* first identifies waypoints after pre-processing and then uses a diffusion map to compute diffusion pseudotime on each cell (Setty et al., 2019). These waypoints are subsequently employed to construct an absorbing Markov chain, which calculates transition probabilities. Finally, pseudotime and branch probabilities derived from the waypoints are projected onto individual cells.

We utilised the *destiny* package to calculate the diffusion map and pseudotime. Subsequent processes—including waypoint selection, absorbing Markov chain construction, probability calculation, and projection—were implemented independently and consolidated into a function called *dandelionR::markovProbability* (**Fig. 3**).

**Fig. 3.**
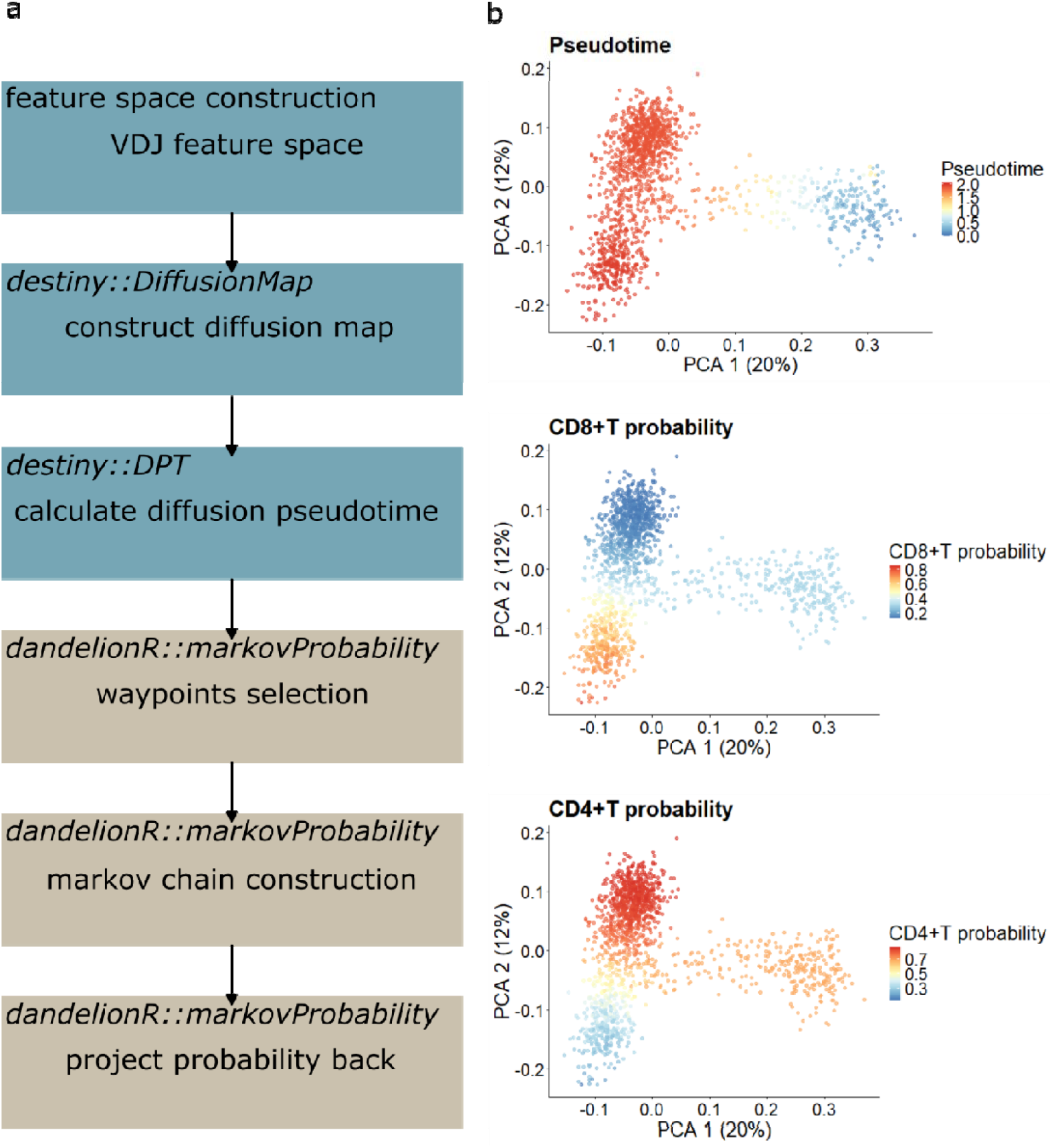
Trajectory analysis with Markov chain in *dandelionR*. (a) *dandelionR*’s trajectory analysis workflow that incorporates outputs from diffusion maps generated by destiny. (b) Pseudotime and branching probabilities of each pseudobulks after trajectory analysis.

Finally, the probabilities and pseudotime of each pseudobulk computed by VDJ feature space were projected back to individual cells through the *dandelionR::projectPseudotimeToCell* function (**Fig. 4**). A total of 39 cells were removed due to not belonging to any neighbourhoods.

**Fig. 4.**
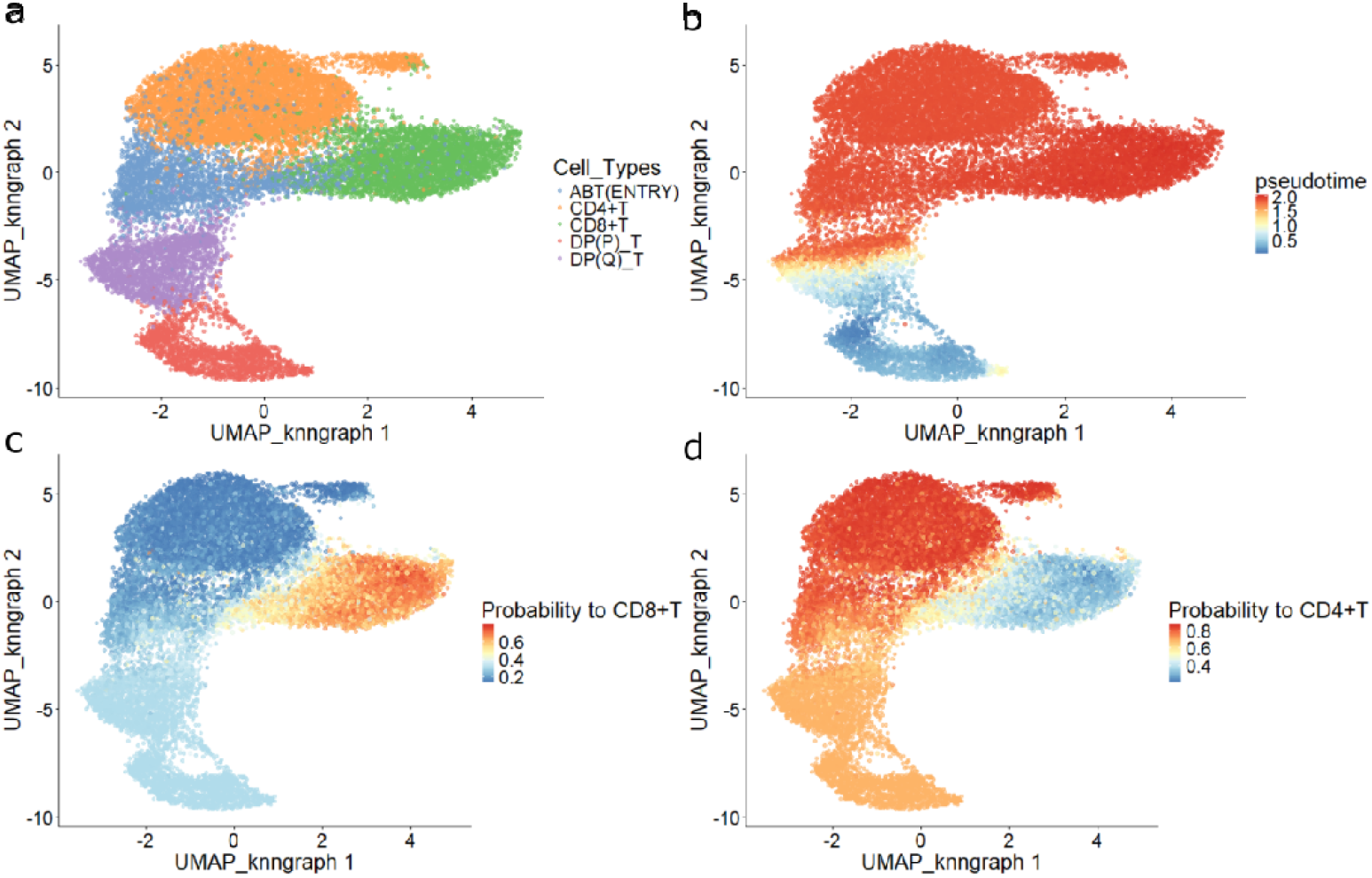
Projection of pseudobulked trajectory results to single cells. (a) Single-cell UMAP plot coloured by cell types. (b, c, d) UMAP coloured by pseudotime, branching probabilities to CD8+T and CD4+T of individual cells.

### Integration with scRepertoire workflow

*scRepertoire* integrates VDJ information with gene expression, storing it in the *colData* of a *SingleCellExperiment* object. The VDJ usage we need lies in column *‘CTgene’*, where V, D, J genes are separated by periods and TRA and TRB chains are separated by an underscore. For instance, the entry *‘TRAV23*.*TRAJ21*.*TRAC_TRBV5-1*.*NA*.*TRBJ2-1*.*TRBC2’* indicates that the TRA chain has V gene TRAV23 and J gene TRAJ21, while TRB chain contains V gene TRBV5-1, unclear D gene, and J gene TRBJ2-1. If any of the chains is absent, it is represented as *‘NA’*, such as *‘TRAV13-2*.*TRAJ23*.*TRAC_NA’*, where the TRB chain is absent.

However, the calculation of VDJ feature space treats each V, D, and J gene separately. To address this, we developed an internal function that extracts the VDJ information from the *‘CTgene’* column generated by *scRepertoire*, splitting the V, D and J genes and storing them in individual columns. This function is incorporated in *dandelionR::setupVdjPseudobulk* to ensure compatibility with *scRepertoire*. Additionally, we introduced parameters to allow users to skip the additional filtering, as it is already performed within *scRepertoire*. To prevent errors from splitting the V, D, J genes correctly, users should first ensure that cells with multiple contigs have already been filtered by setting *‘filterMulti = TRUE’* during *scRepertoire::combineTCR* step.

Overall, the implementations above allowed *dandelionR* to function as a downstream tool for *scRepertoire* (**Fig. 5**), enabling further trajectory analysis after the combination of VDJ information with gene expression data.

**Fig. 5.**
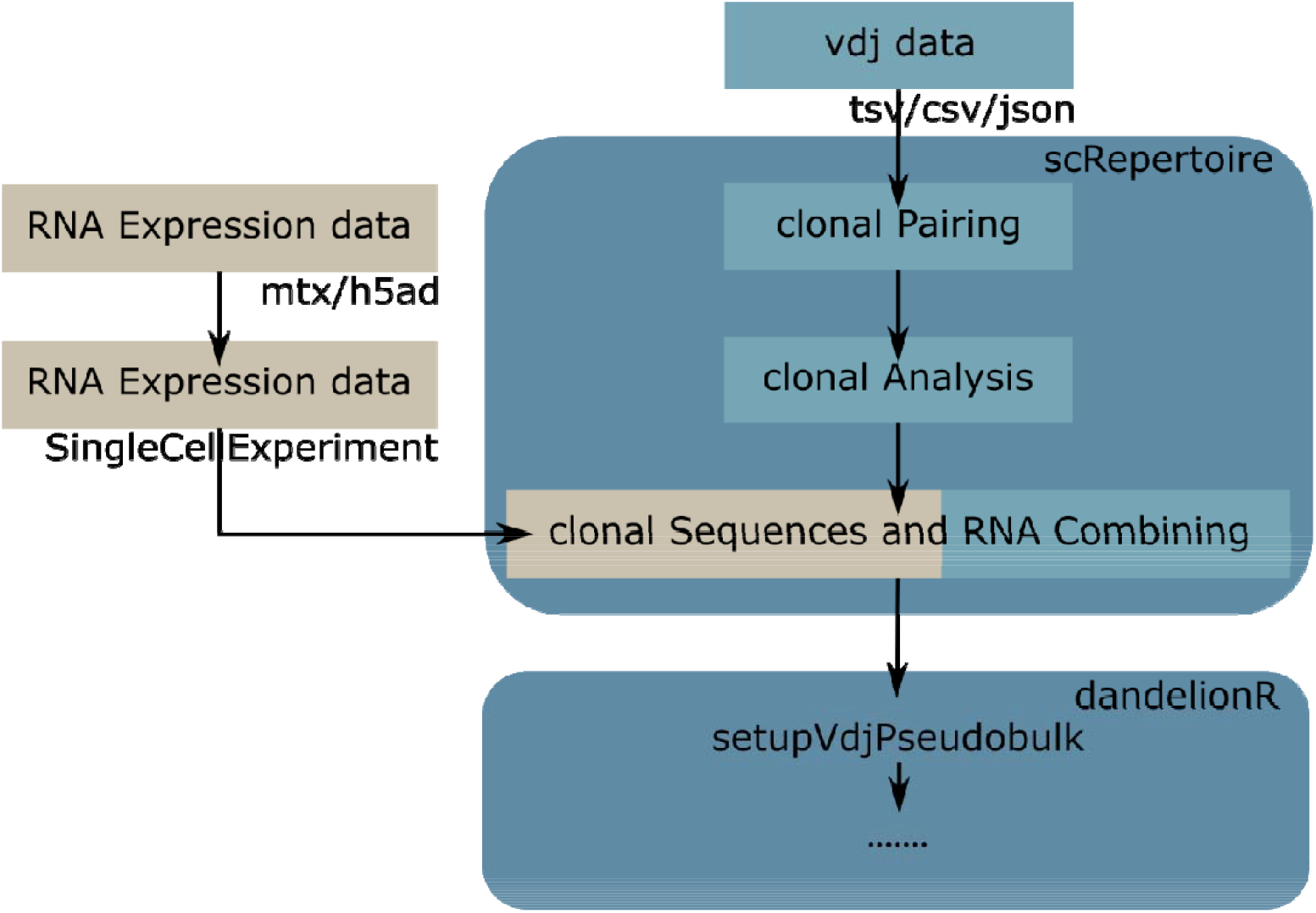
*dandelionR*’s integration with *scRepertoire*’s workflow. Users would perform pre-processing of the scVDJ-seq data as per standard *scRepertoire* workflow and this can then serve as input for *dandelionR*.

## Discussion

We have successfully reproduced *Dandelion*’s trajectory analysis workflow, and incorporated it to *scRepertoire*’s workflow. These steps enable *dandelionR* to function as an R-based trajectory analysis tool that utilised both gene expression data and VDJ information combined by *scRepertoire*.

*dandelionR* implements part of *Palantir*’s trajectory analysis functions based on absorbing Markov chain. Originally, absorbing Markov chain allows the *Palantir* package to handle data with one to multiple terminal states. However, the package we utilised to perform diffusion map, *destiny*, is only suitable for a dataset with a bifurcation topology, which limits the versatility of *dandelionR*. Moreover, *dandelionR* has not yet been applied to B cell development data, which involves more complex processes than T cell development. For instance, after V(D)J recombination, if an immature B cell is self-active, it may re-upregulate RAG to undergo additional rearrangements, a process known as receptor editing (Gay et al., 1993; Okoreeh et al., 2022; Tiegs et al., 1993).

Based on the aforementioned limitations, future directions will focus on replacing *destiny* with alternative methods for conducting diffusion maps and, more importantly, applying *dandelionR* to other real datasets to assess its applicability, evaluate its performance, and explore relevant biological questions.

## Author Contributions

ZKT conceived the project. JY, NB and ZKT wrote the code. JY, and XX performed data analysis. JY and ZKT wrote and edited the manuscript. ZKT supervised the work.

## Acknowledgements

We acknowledge the Children’s Hospital Foundation’s philanthropic contributions awarded to the Ian Frazer Centre for Children’s Immunotherapy Research.

## Reference

Angerer, P., Haghverdi, L., Büttner, M., Theis, F. J., Marr, C., & Buettner, F. (2016). destiny: diffusion maps for large-scale single-cell data in R. Bioinformatics (Oxford, England), 32(8), 1241–1243. 10.1093/bioinformatics/btv715

Borcherding, N., Bormann, N. L., & Kraus, G. (2020). scRepertoire: An R-based toolkit for single-cell immune receptor analysis. F1000Research, 9, 47. 10.12688/f1000research.22139.2

Campbell, K. R., & Yau, C. (2019). A descriptive marker gene approach to single-cell pseudotime inference. Bioinformatics (Oxford, England), 35(1), 28–35. 10.1093/bioinformatics/bty498

Carmona, L. M., & Schatz, D. G. (2017). New insights into the evolutionary origins of the recombination-activating gene proteins and V(D)J recombination. The FEBS Journal, 284(11), 1590–1605. 10.1111/febs.13990

Dann, E., Henderson, N. C., Teichmann, S. A., Morgan, M. D., & Marioni, J. C. (2022). Differential abundance testing on single-cell data using k-nearest neighbor graphs. Nature Biotechnology, 40(2), 245–253. 10.1038/s41587-021-01033-z

Davis, M. M., & Bjorkman, P. J. (1988). T-cell antigen receptor genes and T-cell recognition. Nature, 334(6181), 395–402. 10.1038/334395a0

Deconinck, L., Cannoodt, R., Saelens, W., Deplancke, B., & Saeys, Y. (2021). Recent advances in trajectory inference from single-cell omics data. Current Opinion in Systems Biology, 27(100344), 100344. 10.1016/j.coisb.2021.05.005

Gay, D., Saunders, T., Camper, S., & Weigert, M. (1993). Receptor editing: an approach by autoreactive B cells to escape tolerance. The Journal of Experimental Medicine, 177(4), 999–1008. 10.1084/jem.177.4.999

Heumos, L., Schaar, A. C., Lance, C., Litinetskaya, A., Drost, F., Zappia, L., Lücken, M. D., Strobl, D. C., Henao, J., Curion, F., Single-cell Best Practices Consortium, Schiller, H. B., & Theis, F. J. (2023). Best practices for single-cell analysis across modalities. Nature Reviews. Genetics, 24(8), 550–572. 10.1038/s41576-023-00586-w

Irac, S. E., Soon, M. S. F., Borcherding, N., & Tuong, Z. K. (2024). Single-cell immune repertoire analysis. Nature Methods, 1–16. 10.1038/s41592-024-02243-4

Ji, Z., & Ji, H. (2016). TSCAN: Pseudo-time reconstruction and evaluation in single-cell RNA-seq analysis. Nucleic Acids Research, 44(13), e117. 10.1093/nar/gkw430

Ji, Z., & Ji, H. (2019). Pseudotime reconstruction using TSCAN. Methods in Molecular Biology (Clifton, N.J.), 1935, 115–124. 10.1007/978-1-4939-9057-3_8

Okoreeh, M. K., Kennedy, D. E., Emmanuel, A. O., Veselits, M., Moshin, A., Ladd, R. H., Erickson, S., McLean, K. C., Madrigal, B., Nemazee, D., Maienschein-Cline, M., Mandal, M., & Clark, M. R. (2022). Asymmetrical forward and reverse developmental trajectories determine molecular programs of B cell antigen receptor editing. Science Immunology, 7(74), eabm1664. 10.1126/sciimmunol.abm1664

Saelens, W., Cannoodt, R., Todorov, H., & Saeys, Y. (2019). A comparison of single-cell trajectory inference methods. Nature Biotechnology, 37(5), 547–554. 10.1038/s41587-019-0071-9

Scott, J. K., & Breden, F. (2020). The adaptive immune receptor repertoire community as a model for FAIR stewardship of big immunology data. Current Opinion in Systems Biology, 24, 71–77. 10.1016/j.coisb.2020.10.001

Setty, M., Kiseliovas, V., Levine, J., Gayoso, A., Mazutis, L., & Pe’er, D. (2019). Characterization of cell fate probabilities in single-cell data with Palantir. Nature Biotechnology, 37(4), 451–460. 10.1038/s41587-019-0068-4

Stephenson, E., Reynolds, G., Botting, R. A., Calero-Nieto, F. J., Morgan, M. D., Tuong, Z. K., Bach, K., Sungnak, W., Worlock, K. B., Yoshida, M., Kumasaka, N., Kania, K., Engelbert, J., Olabi, B., Spegarova, J. S., Wilson, N. K., Mende, N., Jardine, L., Gardner, L. C. S., … Haniffa, M. (2021). Single-cell multi-omics analysis of the immune response in COVID-19. Nature Medicine, 27(5), 904–916. 10.1038/s41591-021-01329-2

Street, K., Risso, D., Fletcher, R. B., Das, D., Ngai, J., Yosef, N., Purdom, E., & Dudoit, S. (2018). Slingshot: cell lineage and pseudotime inference for single-cell transcriptomics. BMC Genomics, 19(1), 477. 10.1186/s12864-018-4772-0

Sturm, G., Szabo, T., Fotakis, G., Haider, M., Rieder, D., Trajanoski, Z., & Finotello, F. (2020). Scirpy: a Scanpy extension for analyzing single-cell T-cell receptor-sequencing data. Bioinformatics, 36(18), 4817–4818. 10.1093/bioinformatics/btaa611

Suo, C., Dann, E., Goh, I., Jardine, L., Kleshchevnikov, V., Park, J.-E., Botting, R. A., Stephenson, E., Engelbert, J., Tuong, Z. K., Polanski, K., Yayon, N., Xu, C., Suchanek, O., Elmentaite, R., Domínguez Conde, C., He, P., Pritchard, S., Miah, M., … Teichmann, S. A. (2022). Mapping the developing human immune system across organs. Science (New York, N.Y.), 376(6597), eabo0510. 10.1126/science.abo0510

Suo, C., Polanski, K., Dann, E., Lindeboom, R. G. H., Vilarrasa-Blasi, R., Vento-Tormo, R., Haniffa, M., Meyer, K. B., Dratva, L. M., Tuong, Z. K., Clatworthy, M. R., & Teichmann, S. A. (2024). Dandelion uses the single-cell adaptive immune receptor repertoire to explore lymphocyte developmental origins. Nature Biotechnology, 42(1), 40–51. 10.1038/s41587-023-01734-7

Tiegs, S. L., Russell, D. M., & Nemazee, D. (1993). Receptor editing in self-reactive bone marrow B cells. The Journal of Experimental Medicine, 177(4), 1009–1020. 10.1084/jem.177.4.1009

Tonegawa, S. (1983). Somatic generation of antibody diversity. Nature, 302(5909), 575–581. 10.1038/302575a0

Yang, Q., Safina, K. R., & Borcherding, N. (2024). ScRepertoire 2: Enhanced and efficient toolkit for single-cell immune profiling. In bioRxiv (p. 2024.12.31.630854). 10.1101/2024.12.31.630854

